# The 2017 plague outbreak in Madagascar: data descriptions and epidemic modelling

**DOI:** 10.1101/247569

**Authors:** Van Kinh Nguyen, Cesar Parra-Rojas, Esteban A. Hernandez-Vargas

## Abstract

From August to November 2017, Madagascar endured an outbreak of plague. A total of 2417 cases of plague were confirmed, causing a death toll of 209. Public health intervention efforts were introduced and successfully stopped the epidemic at the end of November. The plague, however, is endemic in the region and occurs annually, posing the risk of future outbreaks. To understand on the plague transmission, we collected real-time data from official reports, described the outbreak’s characteristics, and estimated transmission parameters using statistical and mathematical models. The pneumonic plague epidemic curve exhibited multiple peaks, coinciding with sporadic introductions of new bubonic cases. Optimal climate conditions for rat flea to flourish were observed during the epidemic. Estimate of the plague basic reproduction number *during the large wave of the epidemic* was high, ranging from 5–7 depending on model assumptions. The incubation and infection period for bubonic and pneumonic plague were 4.3 and 3.4 days and 3.8 and 2.9, respectively. Parameter estimation suggested that even with a small fraction of the population exposed to infected rat fleas (1/10000) and a small probability of transition from a bubonic case to a secondary pneumonic case (3%), the high human-to-human transmission rate can still generate a large outbreak. Controlling rodent and fleas can prevent new index cases, but managing human-to-human transmission is key to prevent large-scale outbreaks.

## 1. Introduction

One of the deadliest natural disasters in human history was reported as the Black Death — attributed to the bacterium *Yersinia pestis* — killing about 50 to 200 million people in the 14th century [1]. Although plague was naturally 5 widespread in ancient times, plague outbreaks occurred following the deliberate use and propagation of this disease, serving as a bioweapon [2]. Nowadays, plague epidemics continue to pose a threat to humans, reporting continuous annual occurrence in five countries: Madagascar, Tanzania, Vietnam, China, and the USA [1, 3]. This lethal bacterium can derive in several forms of plague maintaining its existence in a cycle involving rodents and their fleas [4]. While sanitation and public health surveillance have greatly reduced the likelihood of a plague pandemic, isolated plague outbreaks are lethal threats to humankind.

The disease manifests in different clinical forms of plague: bubonic, pneumonic, and septicemic [1]. Human infection is primary driven by bubonic plague, as a result of being bitten by infected fleas. Additionally, direct contamination with infective material can be an alternative transmission route [1]. Patients with bubonic plague can develop sudden onset of fever, headache, chills, tender and painful lymph nodes [5]. While plague can be successfully treated with antibiotics, if untreated, the bacteria can disseminate from the lymph nodes into the bloodstream causing secondary septicemic plague. In addition to the symptoms presented in the bubonic plague, patients with septicemic plague undergo abdominal pain and possibly bleeding into the skin and other organs, at the same time skin and other tissues may turn black and die, especially on fingers, toes, and the nose [4]. However, the most fulminant form of the disease is driven by pneumonic plague that is the only form of plague that can spread from person to person by infectious droplets. The incubation period of primary pneumonic plague is shorter than in the other forms of the disease with an average of 4 days [6]. The disease progresses rapidly and is nearly always fatal without prompt antibiotic treatments [7].

Patients with pneumonic plague often do not transmit the disease to anyone, but in the right conditions one can infect many people and cause an outbreak [7, 8]. This was observed in the last epidemic in Madagascar when an 31-year-old man travelled from the central highlands to the eastern city of Toamasina via the capital city, Antananarivo. He died in transit and dozens of his contacts subsequently became ill [9]. Since then cases of suspected plague have been reported from many areas of Madagascar. On 13 September 2017, the Madagascar Ministry of Public Health notified WHO of an outbreak of pneumonic plague [5]. A total of 2417 cases of plague has been confirmed of which 77% were pneumonic plague, causing until now a death toll of 209 [9]. The Government of Madagascar with the supports of WHO and partners had focused their efforts on strengthening the identification and treatment of patients and their contacts, increasing control of rodents and fleas, and practising safe and dignified burials [10]. These measures have been preventing new cases and deaths, however, the disease is endemic and occurs annually in the region and elsewhere [1, 3], posing the risk of future outbreaks. This paper provides descriptive and numerical analyses of the plague outbreak to facilitate further studies in evaluating the spread of the plague as well as targets for disease control and prevention.

## 2. Materials and Methods

*Outbreak Data - Cumulative Cases.* Data were manually inputted from separate reports of WHO [9], including the cumulative total numbers of clinical cases (confirmed, probable, and suspected). The data can be found at the following link 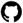 systemsmedicine/plague2017/Cumulative

*Outbreak Data - by Disease Forms.* Data were digitized from the figure reported from WHO [9], including the incidences classified by the three forms of the plague disease: pneumonic, bubonic, and septicemic. The data can be found at the following link 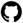 systemsmedicine/plague2017/Classification and the digitized figure can also be found at the same repository.

*Temperature and Precipitation Data.* Data were requested from the National Centers for Environmental Information (Order #1133340 Custom GHCN-Daily CSV). The data can be found at 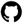 systemsmedicine/plague2017/Climate.

*Descriptive analyses.* With the aim of facilitating modelling works, we described dynamics and patterns of variables that have previously shown to be relevant to plague outbreaks, including temperature and precipitation [11].

*Statistical estimate of the reproduction number.* We estimated the reproduction number (R0) of *Yersina pestitis* using data of pneumonic cases (excluding the bubonic cases) during the second (large) wave of the epidemic, i.e., visually defined from 22/09/17 onwards. The serial interval of plague was assumed gamma distributed with shape and scale parameters are 5.4 and 0.9, respectively [12]. We reported the R0 estimates using several methods for comparison purposes: exponential growth (EG) [13], maximum likelihood (ML) [14], sequential bayesian estimation (SB) [15], binomial assumption [16], and sub-exponential growth model [17, 18]. Implementations in R of the last two methods are available at 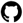 systemsmedicine/plague2017/

*Plague transmission model (PTM).* We proposed a plague transmission model using a modified SEIR (Susceptible-Exposed-Infectious-Removed) model with two additional incorporated components reflecting the seasonal infection from infected rat fleas and imperfect sigmoidal effects of public health interventions. A schematic illustration of the PTM is shown in Figure 1. The model equations are as follows:

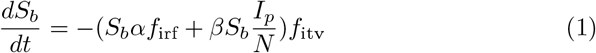

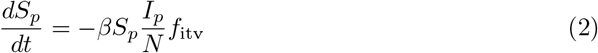

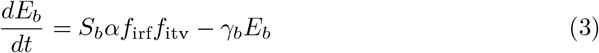

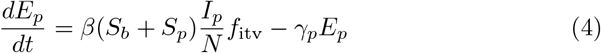

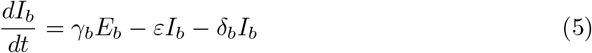

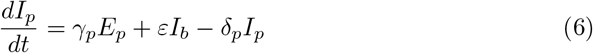

where *S, E, I* describe Susceptible, Exposed, and Infectious and the subscripts *b* and *p* denote bubonic and pneumonic form, respectively. The model assumes that in a population of size *N*, only a small part *S_b_ = pN* is exposed to infected rat fleas. The infected bubonic cases become infectious with a proportion *ε* progressing to pneumonic stage [8]. The overall transmission rates of the two modes flea-to-human and human-to-human are denoted by *α* and *β*, respectively.

**Figure 1:**
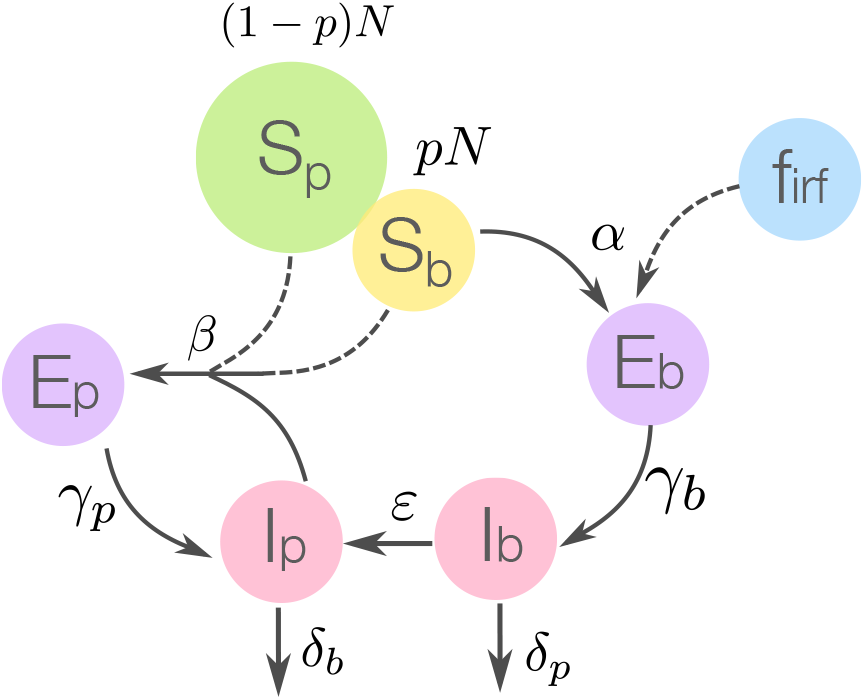
Schematic of plague transmission model (PTM). Assuming a small proportion (p) of the population is exposed to the risk of being bitten by infected rat fleas. The flea density (*f*_irf_) is approximated with a sinusoidal function fitted on Madagascar temperature (see more details in the text of Materials and Methods section). Dashed lines indicate parameters which public health interventions would affect.

It has been shown that climatic conditions favour the survival and reproduction of fleas [19, 20, 21]. Fleas density fluctuates by season temperatures, and changes in their host (rats) also depend on the season which relates to the breeding patterns of rats [22]. To this end, we assumed that the density of infected rat fleas can be described as a sinusoidal function *f*_irf_ = *A + B* sin(2*π*/12*t*) + *C* cos(2*π*/12*t*) [23] following the temperature fluctuation; here the average temperature of Madagascar in the period 1960-2008 [11] was used. As such, the overall flea-to-human transmission parameter *α* implicitly incorporates a scaling factor from temperature to rat fleas density.

We assumed interventions would reduce flea-to-human and human-to-human transmission rates via rodent and flea control, via active case finding and the identification and prophylaxis of contacts. The interventions effect is assumed *imperfect* and has a logistic form as *f*_itv_ = 1 – 1/[1 + *θ* + exp(*τ_h,f_* – *t*)], where *τ* > 0 denotes the time at which the interventions reached half maximum effect in controlling human-to-human (*τ_h_*) and flea-to-human (*τ_f_*) transmission. The reduction from a perfect intervention is defined by *θ* ≥ 0, where *θ* = 0 means a perfect scenario. The infected cases are assumed to recover and die with the total rate of removal from the infected pool being *δ_b_* and *δ_p_* for bubonic and pneumonic cases, respectively.

We fitted the PTM to the daily data of pneumonic and bubonic cases during the large wave of the epidemic curve which was visually defined from 22/09/17 onwards. Model parameters were estimated using the global optimisation algorithm Differential Evolution [24]. We derived also R0 using the next generation matrix [25]. Simulations and estimations were written in R using packages base [26], deSolve [27], and R0 [28] and in Python. Stochastic simulations were performed using a tau-leaping algorithm with a fixed time-step of ca. 15 minutes; code and data are publicly available at 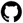 systemsmedicine/plague2017/.

## 3. Results

### Descriptive analyses

During August, bubonic cases appeared sporadically with almost no records of pneumonic form (Fig. 2). An increase in number of pneumonic cases was not necessarily preceded by an increase in the number of bubonic cases (Fig. 2). It seemed to be the epidemic curves include several waves of incidence overlapping each other.

**Figure 2:**
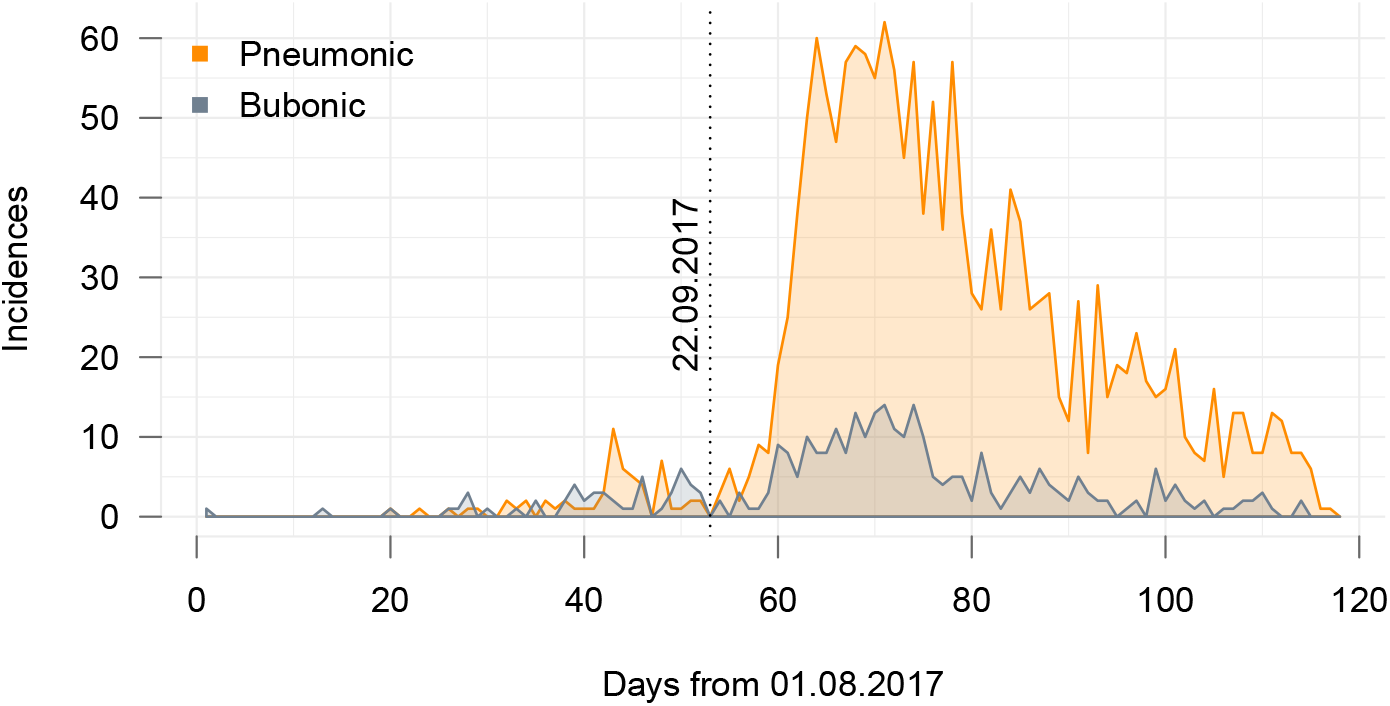
Plague dynamics August-October 2017. Reported incidences of the two forms of plague diseases during the 2017 outbreak. The data were digitized from WHO report’s figure [5]. Data can be accessed and updated by sending merge requests at O smidgroup/plague2017.

Figure 3 shows that plague incidences emerged everyday in the weeks though in some weeks fewer cases were reported during the weekends. No distinctive time lag was observed between the appearances of bubonic and pneumonic cases (Fig. 3). The incidences were negligible during the period when the Famadihana tradition was presumably practised. Precipitation measure exhibited no pattern before or during the outbreak but generally showed a dry climatic condition. Average temperature appeared increasing and reached a higher level (above 23 degree Celsius) around the same time as the outbreak.

**Figure 3:**
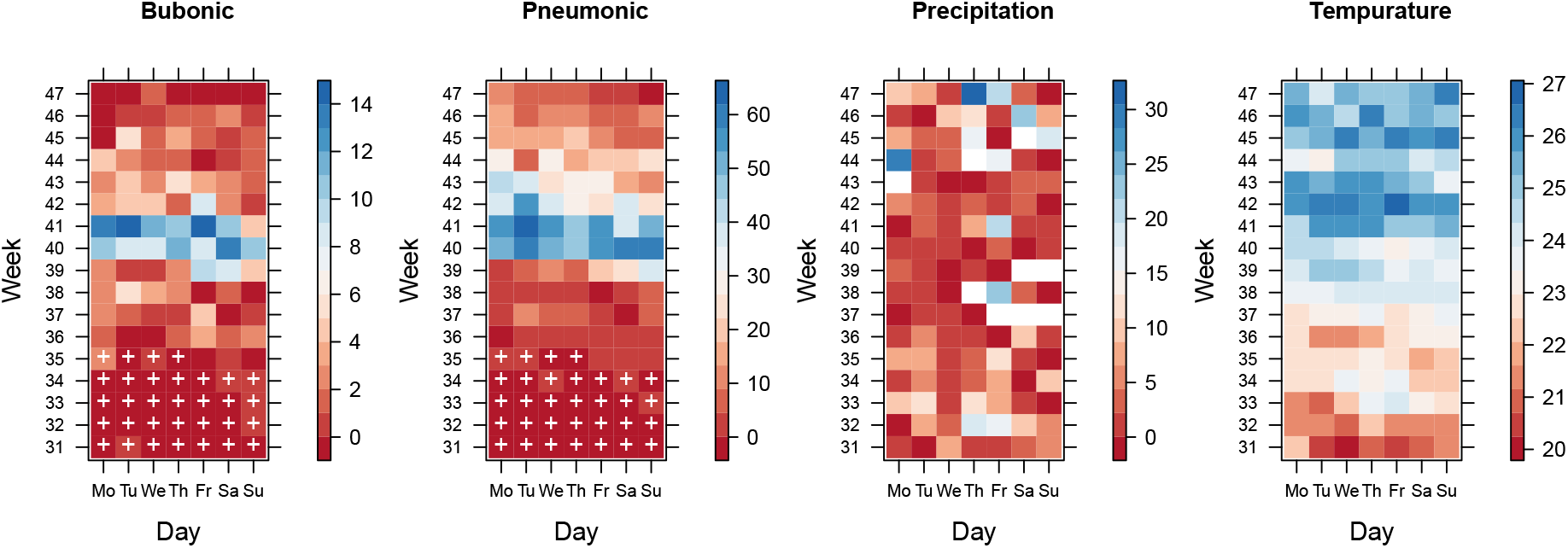
Incidences and climate variables. Reported incidences and average temperature and precipitation classifying by week and day of the weeks. The “+” signs indicate a supposed time period when the Famadihana tradition is practised.

Figure 4 shows that the temperature would typically remain at a level favoring the rat fleas (20–25°C [21]) in the upcoming months and until May in the next year.

**Figure 4:**
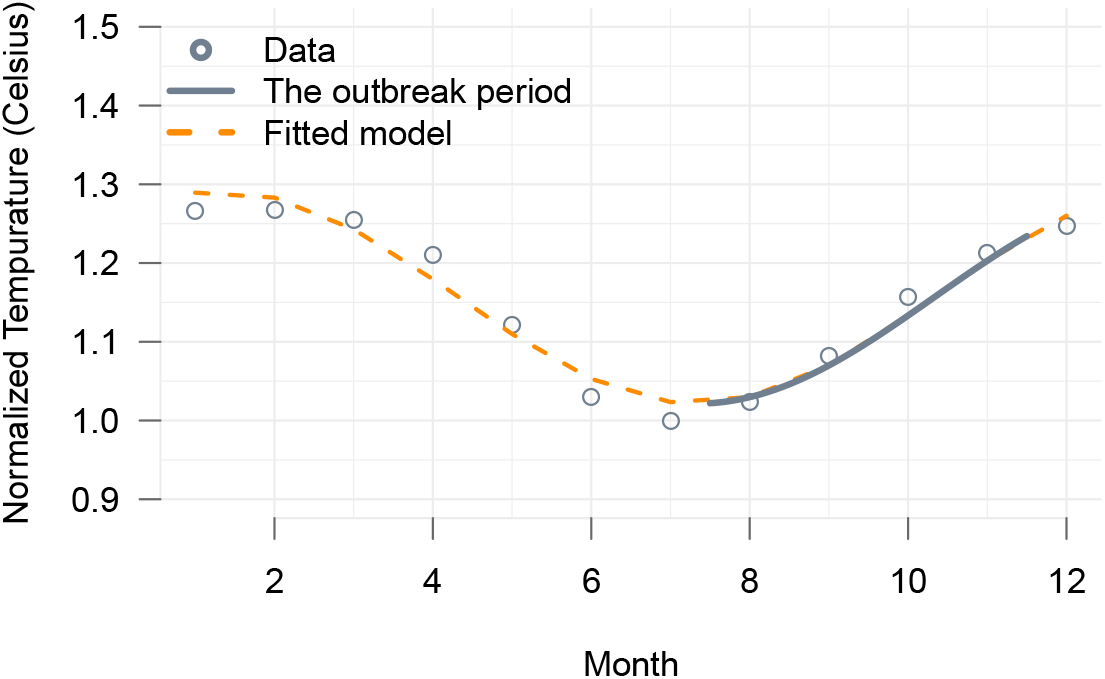
Fitted sinusoidal of Madagascar temperature. Fitted sinusoidal of Madagascar temperature (see Materials and Methods) with *A* = 1.15, *B* = 0.08, *C* = 0.1. The temperature is normalised by the lowest value of July.

### Models and parameter estimates of plague epidemic

Figure 5 shows that the model and the estimated parameters (Table 1) capture well the dynamics of both pneumonic and bubonic data. Stochastic transitions could lead to larger or smaller waves of the dynamics. There are epidemic trajectories that resemble to the intermittent and small epidemic wave during August. The parameters suggest that only a small fraction exposed to infected rat fleas is enough to generate the observed epidemic. Table 1 shows that several statistical estimation methods gave a similar value for plague’s basic reproduction number which is approximately 7. We also estimated R0 using the PTM. Evaluating the Jacobian of the PTM at the disease-free equilibrium yielded a threshold parameter *R*\*

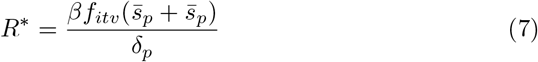

such that *R*\* ≥ 1 results in *λ*_6_ ≥ 0, where *λ*_6_ is the largest eigenvalue of the Jacobian, all the others being non-positive (Appendix 1). This result agrees with the estimate obtained from the next-generation-matrix (NGM) [25], for which we find the leading eigenvalue at *t* = 0 to be 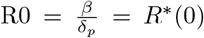. In other words, at the beginning (*S_b_ + S_p_* = *N*) when there are no interventions (*f_itv_* = 1), an infected subject has a transmission rate *β* spent an average 1/*δ_p_* days and thus generate 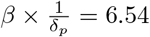 secondary cases.

**Figure 5:**
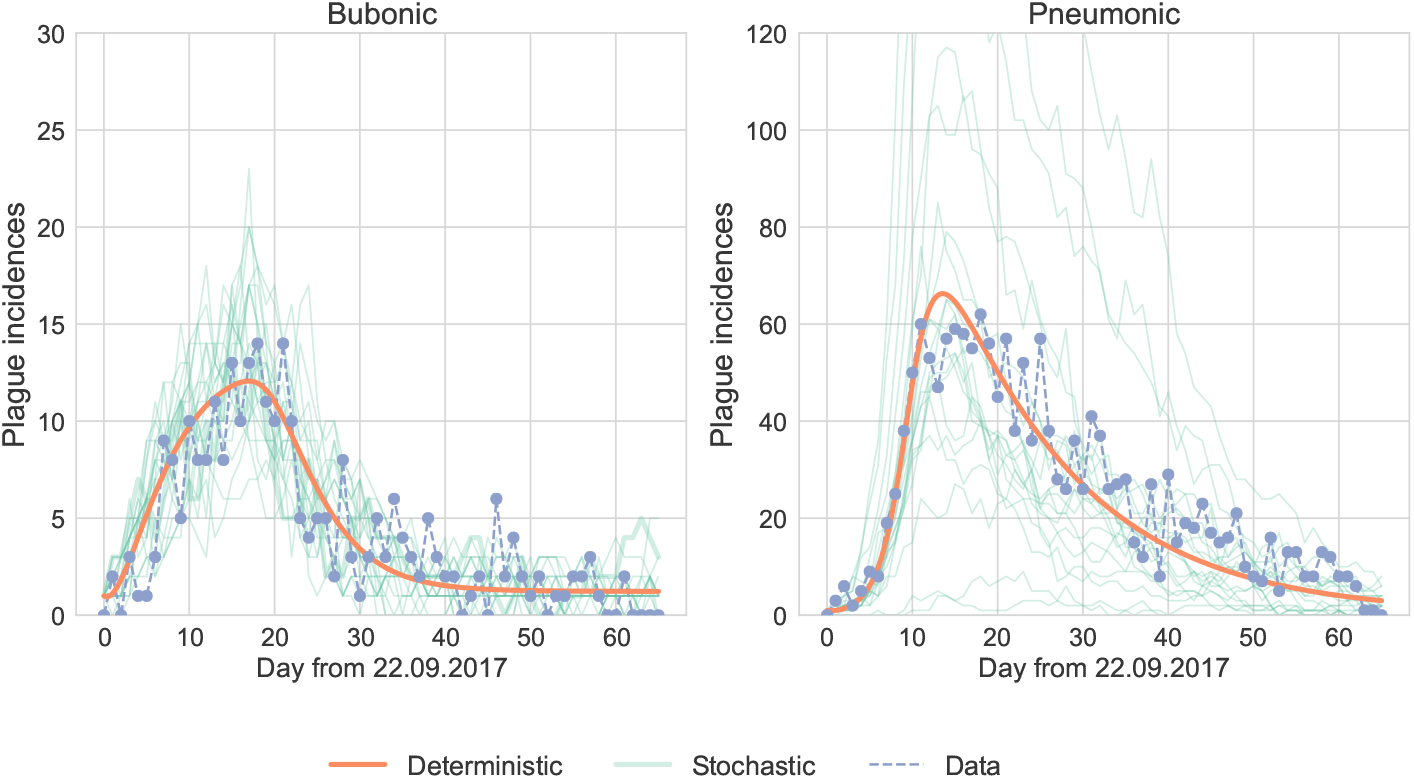
Deterministic and stochastic simulations of plague epidemics. The parameters were estimated with the global optimisation algorithm Differential Evolution. Stochastic simulations were done with tau-leaping algorithm. Python code are publicly available at 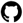 systemsmedicine/plague2017/Fitting.

**Table 1:** Transmission parameters estimated from PTM+ and statistical models.

|  | Estimate | Range | Meaning |
| --- | --- | --- | --- |
| $p$ | $1 \times 10^{-4}$ | 0–0.5 | fraction exposed to infected rat flea |
| $\gamma_b$ | 0.23 | 2–6 days [4] | inverse bubonic plague incubation period |
| $\gamma_p$ | 0.29 | 1–6[8, 7], –12[6] days | inverse pneumonic plague incubation period |
| $\delta_b$ | 0.26 | 4–10 days [29] | inverse bubonic plague infection period |
| $\delta_p$ | 0.34 | 1–12[6], –15[30] days | inverse pneumonic plague infectious period |
| $\varepsilon$ | 0.03 | < 5% [8], 8–10% [29] | rate of transition to secondary pneumonic |
| $\alpha$ | $1.9 \times 10^{-3}$ | 0–10 | (scaled) flea-to-human transmission rate |
| $\beta$ | 2.23 | 0–10 | human-to-human transmission rate |
| $\theta$ | 0.11 | $\geq 0$ | reduction in intervention effects |
| $\tau_h$ | 8.89 | 0–43<sup>++</sup> | time at which interventions reached half its maximum effect on human-to-human |
| $\tau_f$ | 17.93 | 0–43 | — in flea-to-human |
| R0 | Estimate<sup>*</sup> | Methods |  |
| | 6.54 | Next generation matrix[25] ($R0 = \beta/\delta_p$) | |
|  | 6.9[5.2–9.3] | Exponential Growth[13] |  |
|  | 6.9[5.7–8.3] | Maximum likelihood[31] |  |
| | 6.9[0.8–11.7] | Sequential bayes estimation[15] at $t = 0$ | |
|  | 6.9[5.2–9.7] | Binomial assumption[16] |  |
|  | 7.1, 5.1 | Sub-exponential growth[17, 18] at 1 week, at epidemic peak. |  |

<sup>+</sup>Differential Evolution algorithm was run minimizing the least absolute differences of both the bubonic and pneumonic data weighted by their range. The total population used for simulations is $N = 25570895$ [32]. Initial conditions of the model is $\{Np, N(1-p), 0, 0, 1, 1\}$ . For parameters have been previously estimated elsewhere with large variance, their range is used here as boundary constrains for the optimisation. <sup>++</sup>Maximum simulation time. <sup>\*</sup>Number in brackets are 95% Confident Interval.

## 4. Discussion

Mathematical models of infectious diseases have played a central role in understanding epidemics, providing an effective way of assessing disease transmission as well as evaluating disease control and prevention strategies [33]. Mathematical modeling has proposed new vaccination strategies against influenza infection [34]; supported public health strategies for containing emerging influenza pandemics [35, 36] and for the use of antiretroviral treatment for HIV-infected patients as a preventive measure [37]; reported real-time estimates of Ebola’s R
0 to inform the outbreak situation [38], among others.

However, while it has been noted for the last Ebola outbreak [38], data sharing are still poor and WHO practices of reporting (separate PDF files) are putting constraints on modelling works. In the 2014 Ebola outbreak, for example, most of the modelling studies were done rather late in the process [39]. The solution can be as simple as putting a unique Excel file and update it, or better establishing a central website for all WHO outbreak reports, or alternatively, a data hub to encourage user contributed reports.

From modelling aspects, plague outbreaks can be more challenging because: (1) there is a continuous input of flea-to-human transmission (Fig. 2); this implies the observed epidemic curve can be a mixed of multiple waves of infected cases generated from different index cases. Thus, epidemic evaluations could risk to over- or under-estimating the consequences, e.g. the reproduction number or the end time of epidemics; (2) there are a known seasonal pattern of the plague epidemic [11] for which a direct measure of the rat flea population does not exist; (3) The flea-to-human transmission as well as the transition from bubonic cases to pneumonic cases appeared stochastically driven (Fig. 2) and could be highly affected by interventions.

Here, the epidemic curves showed plague incidences appear sporadically during August (Fig. 2). But then a large increase in pneumonic cases was observed, preceding by only a few bubonic cases. Estimates of the plague reproduction number showed that a high estimate is needed to capture this fast growing phase of the epidemic; the estimate doubled the previous estimates ranging 2.8–3.5 [12]. It can be speculated that superspearders were likely to exist in order to generate the larger number of cases in a short time as observed in the pneumonic epidemic curve. This was a common pattern as described in plague outbreaks in Madagascar 2017 [9] and in the US 1919, 1924, and 1980 [8]. Considering the potential mixed of epidemic waves and outbreak locations in the used data, our estimate of R0 could be overestimated. However, the smaller estimates of R0 of this outbreak ranging 1.1-1.4 [40, 41] could be due to the inclusion of the long sporadic incidences occurred during August which can misjudge the transmission ability of pneumonic plague in optimal conditions from 22.09.2017 onwards (Fig. 2).

Nonetheless, approaches adjusting for the propagated outbreak data are needed to further understand its effect on parameter estimates. For example, how do the data of a non-synchronized and combined epidemic curve affect the reproduction number? Implementations and calculations using current methods raised also an issue that has been discussed elsewhere [31], that is estimating the reproduction number using serial interval could yield very high value during the first days of the epidemic growing phase as the denominator of the estimator is extremely small during this period. Overestimate of R0 has also been observed in populations with heterogeneous contact pattern population [42].

It appeared that sporadic inputs of bubonic cases brought new index cases to the human-to-human transmission network constantly with an estimate of transition to secondary pneumonic is of 3% (Fig. 2). It followed that the pneu monic epidemic curve exhibited multiple peaks and waves. As the transmission rate can be high, this observation suggest that vector control are key to prevent potential next waves of the epidemic. This might be practical as the fraction of population exposed to infected rat fleas was low (Table 1). On the other hand, the estimate of human-to-human transmission needs to be large in order to capture the data. This prompts that intervention efforts, in cases of exhaustible of resources, should prioritise stopping human-to-human transmission route. In this case, while the bubonic cases would continue to appear they would not able to generate large size outbreaks. This is practical as with the usual public health intervention of early detection of the incidences would not only stop the human-to-human transmission but also provide a better chance for new bubonic cases to be treated early.

Experiments have shown that an optimal climate for rat fleas to flourish is a dry climate with temperatures of 20–25°C [21]. These conditions were observed during the outbreak: a generally dry weather with the optimal temperature coincided with the period of high epidemic activity. As further shown in Fig. 4, these conditions would typically remain the same until next May, see further in Kreppel et al. [11]. This again stresses the role of vector control in preventing the next waves. It is worth noting that the global changes in climate conditions could pose a risk of irregular changes in vector biting rate and reproduction [43], requiring affected regions to be vigilant in plague control and prevention. For the modelling aspect, studies on the dynamics of rat fleas are needed to help further parameterise the plague transmission model.

In this paper, we collected and described relevant data of the 2017 plague epidemic in Madagascar. We proposed a working mathematical model for evaluating and predicting epidemic consequences and what-if scenarios. We discussed potential drawbacks in modelling propagated epidemic data. We hope that the results would contribute informative insights for public health officers and provide a framework for further understanding the dynamics of plague outbreaks.

## Authors contributions

VKN, EAHV: conceptualization and data curation. VKN: methodology, formal analysis, investigation, and visualization. CPR: formal analysis, stochastic simulations implementation, and visualization. EAHV: funding acquisition, supervision. All authors: review & editing.

## Acknowledgment

This work was supported by the Alfons und Gertrud Kassel-Stiftung.

## Supporting Information

The eigenvalues of the Jacobian of the PTM evaluated at the disease-free equilibrium 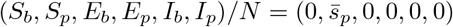 have the form

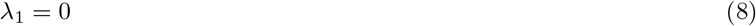

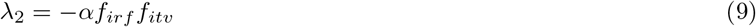

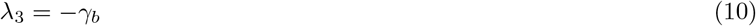

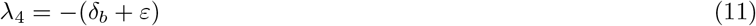

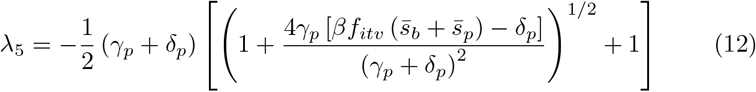

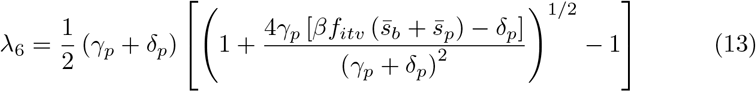

## References

[1] N. A. Boire, V. A. A. Riedel, N. M. Parrish, S. Riedel, Lessons Learned from Historic Plague Epidemics: The Relevance of an Ancient Disease in Modern Times, Journal of Infectious Diseases & Preventive Medicine 2 (2). doi:10.4172/2329-8731.1000114.

[2] S. Riedel, Plague: from natural disease to bioterrorism., Proceedings (Baylor University. Medical Center) 18 (2) (2005) 116–124.

[3] D. T. Dennis, K. L. Gage, N. G. Gratz, J. D. Poland, E. Tikhomirov, World Health Organization. Epidemic Disease Control, Plague manual: epidemiology, distribution, surveillance and control.

[4] Centers for Disease Control and Prevention. Plague [online] (2005).

[5] World Health Organization, Plague – Madagascar, WHO. URL http://www.who.int/csr/don/02-november-2017-plague-madagascar/en/

[6] S. L. Raymond Gani, Epidemiologic Determinants for Modeling Pneumonic Plague Outbreaks, Emerging Infectious Diseases 10 (4) (2004) 608–614. doi:10.3201/eid1004. 030509. URL http://wwwnc.cdc.gov/eid/article/10/4/03-0509_article.htm

[7] P. S. Mead, Plague in Madagascar — A Tragic Opportunity for Improving Public Health, New England Journal of Medicine 378 (2) (2017) 106–108. doi:10.1056/NEJMp1713881. URL http://www.nejm.org/doi/10.1056/nejmp1713881

[8] J. L. Kool, R. A. Weinstein, Risk of Person-to-Person Transmission of Pneumonic Plague, Clinical Infectious Diseases 40 (8) (2005) 1166–1172. doi:10.1086/428617. URL https://academic.oup.com/cid/article-lookup/doi/10.1086/428617

[9] World Health Organization, Plague outbreak situation reports.

[10] World Health Organization, WHO | Madagascar’s plague epidemic is slowing, but we must sustain the response, WHO. URL http://www.who.int/mediacentre/news/releases/2017/plague-madagascar-slowing/en/

[11] K. S. Kreppel, C. Caminade, S. Telfer, M. Rajerison, L. Rahalison, A. Morse, M. Baylis, A Non-Stationary Relationship between Global Climate Phenomena and Human Plague Incidence in Madagascar, PLOS Neglected Tropical Diseases 8 (10) (2014) e3155. doi: 10.1371/journal.pntd.0003155. URL http://dx.plos.org/10.1371/journal.pntd.0003155

[12] H. Nishiura, M. Schwehm, M. Kakehashi, M. Eichner, Transmission potential of primary pneumonic plague: time inhomogeneous evaluation based on historical documents of the transmission network, Journal of Epidemiology & Community Health 60 (7) (2006) 640–645. doi:10.1136/jech.2005.042424 http://doi.wiley.com/10.1111/j.1750-2659.2009.00106.x.

[13] M. L. J Wallinga, How generation intervals shape the relationship between growth rates and reproductive numbers, Proceedings of the Royal Society B: Biological Sciences 274 (1609) (2007) 599–604. doi:10.1098/rspb.2006.3754.

[14] L. F. White, J. Wallinga, L. Finelli, C. Reed, S. Riley, M. Lipsitch, M. Pagano, Estimation of the reproductive number and the serial interval in early phase of the 2009 influenza A/H1N1 pandemic in the USA, Influenza and Other Respiratory Viruses 3 (6) (2009) 267–276. doi:10.1111/j.1750-2659.2009.00106.x. URL http://doi.wiley.com/10.1111/j.1750-2659.2009.00106.x

[15] L. M. A. Bettencourt, R. M. Ribeiro, Real Time Bayesian Estimation of the Epidemic Potential of Emerging Infectious Diseases, PLoS ONE 3 (5) (2008) e2185. doi:10.1371/journal.pone.0002185.

[16] H. Nishiura, Correcting the Actual Reproduction Number: A Simple Method to Estimate R0 from Early Epidemic Growth Data, International Journal of Environmental Research and Public Health 7 (1) (2010) 291–302. doi:10.3390/ijerph7010291. URL http://www.mdpi.com/1660-4601/7/1/291

[17] C. Viboud, L. Simonsen, G. Chowell, A generalized-growth model to characterize the early ascending phase of infectious disease outbreaks, Epidemics 15 (2016) 27–37. doi: 10.1016/j.epidem.2016.01.002. URL http://linkinghub.elsevier.com/retrieve/pii/s1755436516000037

[18] G. Chowell, C. Viboud, L. Simonsen, S. M. Moghadas, Characterizing the reproduction number of epidemics with early subexponential growth dynamics, Journal of The Royal Society Interface 13 (123) (2016) 20160659. doi:10.1098/rsif.2016.0659. URL http://rsif.royalsocietypublishing.org/lookup/doi/10.1098/rsif.2016.0659

[19] R. J. Eisen, K. L. Gage, Transmission of Flea-Borne Zoonotic Agents*, dx.doi.org 57 (1) (2011) 61–82. doi:10.1146/annurev-ento-120710-100717. URL http://www.annualreviews.org/doi/10.1146/annurev-ento-120710-100717

[20] K. Chotelersak, C. Apiwathnasorn, S. Sungvornyothin, C. Panasoponkul, Y. Samung, J. Ruangsittichai, Correlation Of Host Specificity, Environmental Factors And Oriental Rat Flea Abundance., The Southeast Asian journal of tropical medicine and public health 46 (2) (2015) 198–206. URL https://www.ncbi.nlm.nih.gov/pubmed/26513922

[21] J. F. D. Shrewsbury, A History of Bubonic Plague in the British Isles, Cambridge University Press, 2005. URL https://books.google.nl/books?id=ATOlhaEvN3wC

[22] N. Varlik, A Natural History of Plague, Cambridge University Press, 2015, p. 17–54. doi:10.1017/CBO9781139004046.002.

[23] J. L. Aron, R. M. May, The population dynamics of malaria, in: The Population Dynamics of Infectious Diseases: Theory and Applications, Springer US, Boston, MA, 1982, pp. 139–179. doi:10.1007/978-1-4899-2901-3_5. URL http://link.springer.com/10.1007/978-1-4899-2901-3_5

[24] K. M. Mullen, D. Ardia, D. L. Gil, D. Windover, J. Cline, DEoptim: An R Package for Global Optimization by Differential Evolution, Journal of Statistical Software 40 (1) (2011) 1–26. doi:10.18637/jss.v040.i06.

[25] O. Diekmann, J. Heesterbeek, M. Roberts, The construction of next-generation matrices for compartmental epidemic models, Journal of the Royal Society Interface (2009) rsif20090386.

[26] R Core Team, R: A Language and Environment for Statistical Computing, R Foundation for Statistical Computing, Vienna, Austria (2017). URL https://www.r-project.org/

[27] K. Soetaert, T. Petzoldt, R. W. Setzer, Solving Differential Equations in R: Package deSolve, Journal of Statistical Software 33 (9). doi:10.18637/jss.v033.i09. URL http://www.jstatsoft.org/v33/i09/

[28] T. Obadia, R. Haneef, P.-Y. Boёlle, The R0 package: a toolbox to estimate reproduction numbers for epidemic outbreaks, BMC Medical Informatics and Decision Making 12 (1) (2012) 212. doi:10.1186/1472-6947-12-147. URL http://bmcmedinformdecismak.biomedcentral.com/articles/10.1186/1472-6947-12-147

[29] V. V. Nikiforov, H. Gao, L. Zhou, A. Anisimov, Plague: Clinics, Diagnosis and Treatment, in: Yersinia pestis: Retrospective and Perspective, Springer, Dordrecht, Dordrecht, 2016, pp. 293–312. doi:10.1007/978-94-024-0890-4_11. URL https://link.springer.com/chapter/10.1007/978-94-024-0890-4_11

[30] T. H. Tieh, E. Landauer, Primary pneumonic plague in Mukden, 1946, and report of 39 cases with three recoveries., Journal of Infectious Diseases 82 (1) (1948) 52–58. URL http://eutils.ncbi.nlm.nih.gov/entrez/eutils/elink.fcgi?dbfrom=pubmed&id=18898004&retmode=ref&cmd=prlinks

[31] L. Forsberg White, M. Pagano, A likelihood-based method for real-time estimation of the serial interval and reproductive number of an epidemic, Statistics in Medicine 27 (16) (2008) 2999–3016. doi:10.1002/sim.3136. URL http://doi.wiley.com/10.1002/sim.3136

[32] United Nations Department of Economic and Social Affairs. World Population Prospects: The 2017 Revision [online] (2017).

[33] H. Heesterbeek, R. M. Anderson, V. Andreasen, S. Bansal, D. De Angelis, C. Dye, K. T. D. Eames, W. J. Edmunds, S. D. W. Frost, S. Funk, T. D. Hollingsworth, T. House, V. Isham, P. Klepac, J. Lessler, J. O. Lloyd-Smith, C. J. E. Metcalf, D. Mollison, L. Pellis, J. R. C. Pulliam, M. G. Roberts, C. Viboud, Isaac Newton Institute IDD Collaboration, Modeling infectious disease dynamics in the complex landscape of global health, Science 347 (6227) (2015) aaa4339–aaa4339. doi:10.1126/science.aaa4339. URL http://www.sciencemag.org/cgi/doi/10.1126/science.aaa4339

[34] M. A. Rose, O. Damm, W. Greiner, M. Knuf, P. Wutzler, J. G. Liese, H. Krüger, U. Wahn, T. Schaberg, M. Schwehm, T. F. Kochmann, M. Eichner, The epidemiological impact of childhood influenza vaccination using live-attenuated influenza vaccine (LAIV) in Germany: predictions of a simulation study, BMC Infectious Diseases 14 (1) (2014) 379. doi:10.1186/1471-2334-14-40. URL http://bmcinfectdis.biomedcentral.com/articles/10.1186/1471-2334-14-40

[35] N. M. Ferguson, D. A. T. Cummings, S. Cauchemez, C. Fraser, S. Riley, A. Meeyai, S. Iamsirithaworn, D. S. Burke, Strategies for containing an emerging influenza pandemic in Southeast Asia. - PubMed - NCBI, Nature 437 (7056) (2005) 209–214. doi:10.1038/ nature04017.

[36] I. M. Longini, Containing pandemic influenza at the source. - PubMed - NCBI, Science 309 (5737) (2005) 1083–1087. doi:10.1126/science.1115717.

[37] F. Tanser, T. Barnighausen, E. Grapsa, J. Zaidi, M. L. Newell, High coverage of ART associated with decline in risk of HIV acquisition in rural KwaZulu-Natal, South Africa. - PubMed - NCBI, Science 339 (6122) (2013) 966–971. doi:10.1126/science.1228160.

[38] C. L. Althaus, Estimating the Reproduction Number of Ebola Virus (EBOV) During the 2014 Outbreak in West Africa, PLOS Currents Outbreaks doi:10.1371/currents. outbreaks.91afb5e0f279e7f29e7056095255b288. URL http://currents.plos.org/outbreaks/?p=40381

[39] J.-P. Chretien, S. Riley, D. B. George, Mathematical modeling of the West Africa Ebola epidemic, eLife 4 (2015) e09186. doi:10.7554/eLife.09186.

[40] S. Tsuzuki, H. Lee, F. Miura, Y. H. Chan, S.-m. Jung, A. R. Akhmetzhanov, H. Nishiura, Dynamics of the pneumonic plague epidemic in Madagascar, August to October 2017, Eurosurveillance 22 (46) (2017) e0005887. doi:10.2807/1560-7917.ES.2017.22.46. 17-00710. URL http://www.eurosurveillance.org/content/10.2807/1560-7917.ES.2017.22.46.17-00710

[41] H. Nishiura, P. Yan, C. K. Sleeman, C. J. Mode, Estimating the transmission potential of supercritical processes based on the final size distribution of minor outbreaks. - PubMed - NCBI, Journal of Theoretical Biology 294 (2012) 48–55. doi:10.1016/j.jtbi.2011.10.039. URL http://linkinghub.elsevier.com/retrieve/pii/s0022519311005625

[42] C. Llensa, D. Juher, J. Saldaña, On the early epidemic dynamics for pairwise models, Journal of Theoretical Biology 352 (2014) 71–81. doi:10.1016/j.jtbi.2014.02.037. URL http://linkinghub.elsevier.com/retrieve/pii/S0022519314001210

[43] World Health Organization, WHO | Climate change and human health - risks and responses. Summary., WHO. URL http://www.who.int/globalchange/climate/summary/en/index5.html

